# Catch me if you can: *Arabidopsis thaliana* lags in adaptation to contemporary climate change

**DOI:** 10.64898/2026.06.02.729671

**Authors:** Laura Leventhal, Moises Exposito-Alonso

## Abstract

Anthropogenic climate change fosters unprecedented temperature challenges, with each year breaking a temperature record. Through evolution by natural selection, species and their populations have adapted to their previously local environments. However, as the average global land temperature has increased by ~2 °C or more, natural selection on many species may not act fast enough with climate change, creating an adaptation lag. To understand potential adaptation lags to recent climate change, we conducted a meta-analysis on the largest set of single-species field transplantation experiments across climates with the broadly-distributed model plant, *Arabidopsis thaliana*, with a total of 1,600 germplasm and 42 field trials. We developed a Gaussian fitness model dependent on local environment and climate deviations to infer genotype-specific adaptation lag parameters. We estimate a mean thermal adaptation lag over 1.91 °C, suggesting that local populations, on average, are better adapted when transplanted to locations cooler than their home climates. While a less than 2 °C temperature mismatch appears small, its impact on fitness corresponds to a 14% cumulative burden over time, which compounds depending on the future climate emission scenario. Combining climate model projections under different scenarios, we found that by 2025, populations would have lost 30% demographic potential under a moderate emissions scenario. Our discovery of this adaptation lag shows that even this highly adaptable species has not kept pace with recent climate change.

---

Global mean temperatures have already increased by ~2 °C, substantially threatening biological diversity (Exposito-Alonso et al., 2022; Habibullah et al., 2022; National Climatic Data Center, 2025). Scientists have previously documented species responses to warming through migration: by shifting their geographic ranges by 6.1 km per decade poleward and 29 m per decade in altitude (Lenoir et al., 2008; Parmesan & Yohe, 2003; Ramírez-Barahona et al., 2025) or through phenological strategies: by advancing the timing of key adaptive life-history events, such as flowering 2.3 days earlier per decade (Parmesan, 2006; Parmesan & Yohe, 2003; Wiens, 2016). However, whether populations are evolving and adapting to the new environmental conditions, or whether they are already mismatched and lagging behind climate change, remains poorly understood. Here we introduce a framework to estimate adaptation lag using large-scale common garden experiments across climates in plants in combination with climate deviations of the last decades for inference.

\ The first studies investigating adaptation to different climates in the early 20^th^ century asked whether populations within a species harbor genetic adaptations to different local environments with the knowledge that plants often experience a range of environments within their distribution (Clausen et al., 1941; Turesson, 1922). Turesson in Sweden, and Clausen, Keck, and Hiesey in California, tested this idea by collecting plants of the same species along altitudinal or latitudinal gradients, and planted them reciprocally in common gardens, quantitatively showing local adaptation: populations harbor local advantage measured by increased fitness in gardens near their locations of origin. This pattern has been replicated in hundreds of species (Hereford, 2009), and combining genome sequencing has confirmed such local adaptation has a genetic basis (Expósito-Alonso et al., 2019; Fournier-Level et al., 2011; Savolainen et al., 2013). Given its prevalence in ecology and evolution research, local adaptation data has accumulated and a recent meta-analysis showed that with just 2 °C of warming, the fitness advantage conferred by local adaptation was lost, functionally erasing the signal of local adaptation (Bontrager et al., 2020), warning that an adaptation lag may have already occurred.

We suspected that an adaptation lag to global warming may be already detectable in some species, although we expect the statistical signal across a species to be weak and challenging to infer, as many species have broad geographic ranges spanning many degrees Celsius in annual average temperature. Still, we aimed to quantify lag in the model plant *Arabidopsis thaliana*, whose geographic range spans most temperate regions across continents, and where common gardens have been deployed across dozens of locations worldwide (Leventhal et al., 2025). In total, we synthesize fitness data from up to 1614 *A. thaliana* populations transplanted across 42 experimental gardens (**Fig. 1, Table S1–S2**). Seeds from most of these populations (i.e. genotypes) were collected in the 90s and early 2000s (although the year of collection range for our data set is 1937-2012) from known GPS locations and maintained in seed stocks. In this meta-analysis, these seed stocks function as pseudo-”resurrection” plants relative to their collection year (Franks et al., 2007, 2018). Because the accessions were grown in controlled environments after collection, they are essentially frozen in evolutionary time since there was a no-selection environment and the accumulation of new mutations is low per generation and likely neutral. These banked genotypes were maintained by several organizations and then were often re-used in multiple common gardens, most of which were conducted during the 2010s-2020s, a time period which experienced substantial climate deviations (**Fig. 1C**, climate TerraClim (Abatzoglou et al., 2018)).

**Figure 1.**
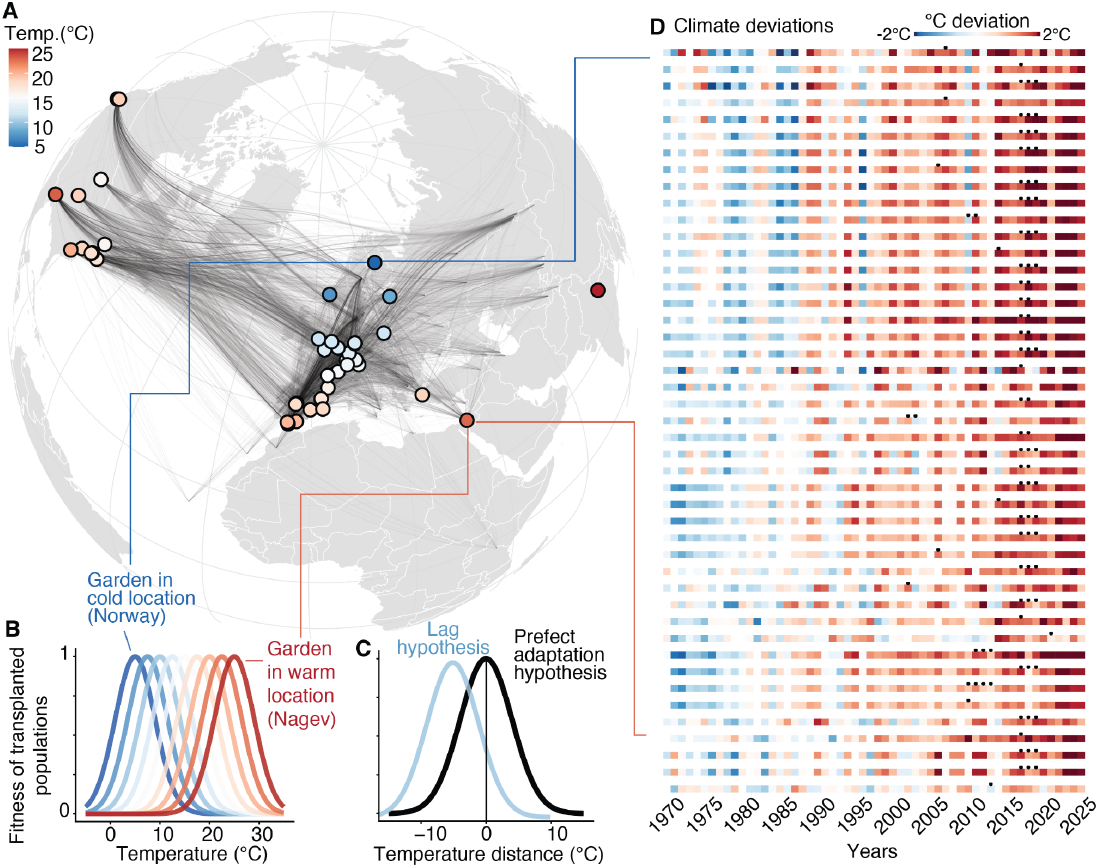
42 transplant common gardens and climate deviations for meta-analysis of adaptation lag. (A) Global projection of common gardens used in this study and the population (or accessions) that were planted in each location. Points represent the garden locations and are colored by temperature, while black lines represent accessions. The lines start at the population’s home coordinates and end in the garden(s) in which they were planted. (B) Representation of local adaptation with Gaussian fitness curves, where the fitness maximum is reached when the temperature of the site matches the home temperature of the genotype. (C) Adaptation lag visualization, where if fitness maxima is reached at no temperature distance, then no adaptation lag occurs. A left shifted fitness peak suggests adaptation lag consistent with climate change predictions, where a genotype is adapted to an environment cooler than their current environment. (D) Diagram of climate deviations by experimental site use in this meta analysis. Gardens are sorted by latitude, and points on the bars represent the year the experiment was conducted in that location. The climate deviations were calculated by subtracting the Terraclim temperature value from the given year by the mean baseline annual temperature for that area (1970-2000). Deviations are all relative to that garden location’s specific baseline.

With this dataset in hand, we built a Gaussian fitness model under an assumption of local adaptation: each population is expected to achieve maximal fitness when the temperature of the common garden matches the temperature of its home site. We define the temperature mismatch as *d = T*_*s*_ – *T*_*g*_, where *T*_*s*_ is the mean annual temperature of the common garden site, *T*_*g*_ is the mean annual temperature at the genotype’s location of origin in the year it was collected. With increasing distance, fitness follows Gaussian function decay, such that fitness is maximized at *d* = 0 and declines as |*d*| increases. Across species, fitness responses to temperature often take a Gaussian form, as there is often a genotype or population specific optimal temperature (Latimer et al., 2015; Wooliver et al., 2022). If populations were lagging behind their optima, then the environmental transplant distance is further increased by a λ lag: *w*_*observed*_ *= w*_*max*_ *exp(−(d* + *λ)*^*2*^*/2V*_*s*_ ); which would be visually a left-shifted Gaussian curve (**Fig. 1C**).

We estimated the parameters of our Gaussian function by transforming this function into a log quadratic regression fitted within a Bayesian hierarchical framework to account for various random effects (see **Methods** and full derivation, symbol definition **Table S3**, model terms **Table S4**, raw fitness data **Fig. S1**, and model diagnostics **Fig. S3–4–5–6)**, with *λ =* −1.91 °C (95% CI: 0.99-3.18) (**Fig. 2A**). The Gaussian curve decay is broad, *V*_*s*_ = 824 (95 % CI [698, 1004]), where *V*_*s*_ is the variance parameter of the Gaussian fitness function (i.e., the squared width of the fitness peak, such that larger values correspond to weaker stabilizing selection and slower fitness decline away from the optimum). This large value indicates a shallow fitness gradient, corroborating the common knowledge that *A. thaliana* has a broad environmental distribution; accordingly, the per-year fitness cost of a 1 °C temperature deviation from the optimum is small (~0.1 %). However, the effect of even a small deviation from the optimum can be catastrophic for a small annual plant in which fitness compounds multiplicatively through generations, because even small per-generation fitness reductions are applied to population growth each generation, leading to cumulative declines over time (**Fig. S7**). This is consistent with patterns shown previously in an *A. thaliana* experiment spanning four locations in Europe, where in the coldest common garden location of Oulu (Finland), accessions several degrees south in latitude had the highest fitness, not the closest local population (Wilczek et al., 2014). Our model showed that individual experiments, inducing the original lag experiment or any single study analyzed independently, did not have sufficient power to detect lag (see drop-one-out and leave-one-in re-analyses in **Supplemental Materials, Table S8**). In contrast, the accumulation of experiments over recent decades, particularly the addition of over 30 locations from a large-scale experimental evolution project, provided the power necessary to infer an adaptation lag.

**Figure 2.**
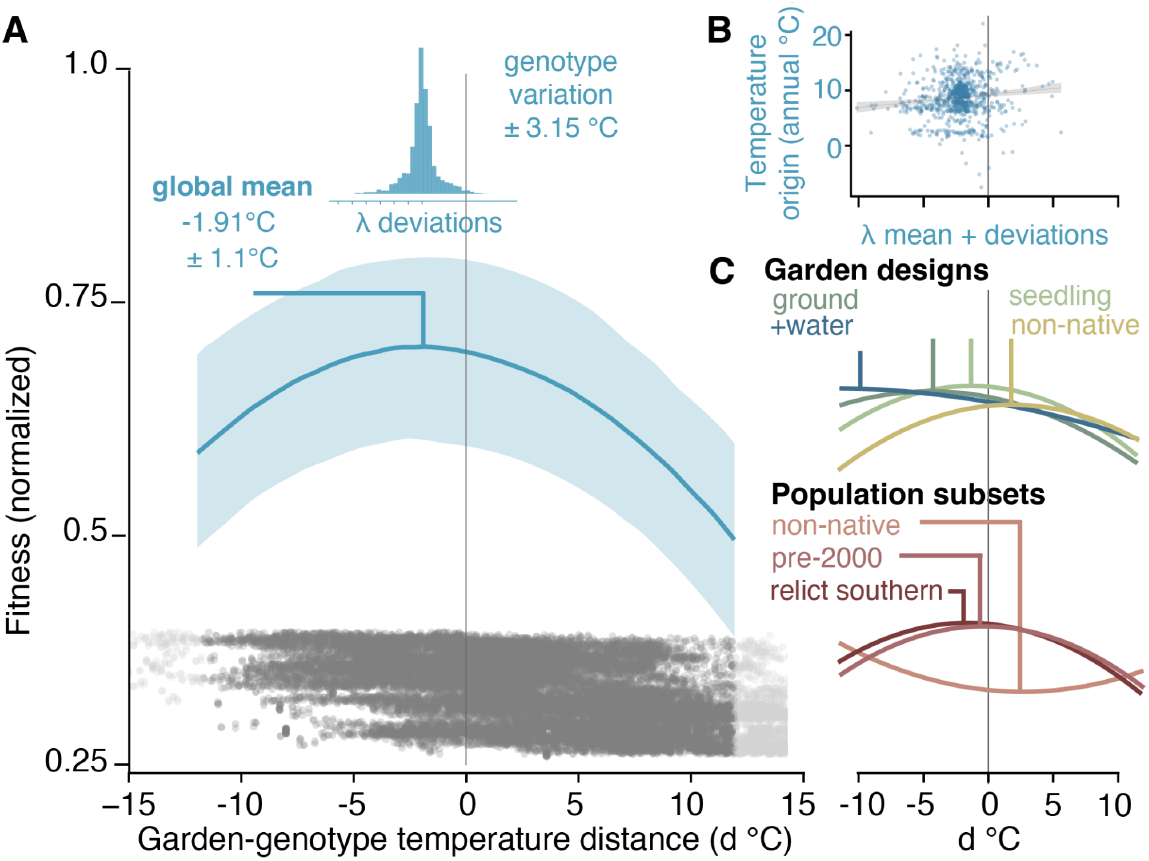
Estimate of global adaptation lag across *Arabidopsis thaliana*. (A) Mean adaptation lag for accessions in our meta analysis is at −1.91 °C, suggesting adaptation to cooler environments. (B) Spread of adaptation lag across genotypes. X-axis is genotype-level adaptation lag minus mean adaptation of all genotypes, showing mean lag therefore centered at zero. (C) Curiosity tests of different subsets of the data on adaptation lag. We grouped our data set by garden design parameters (top) such as if the plants starting as seeds or seedlings outside, were planted in the ground vs. pots, had a water addition treatment, and finally if the garden was located in the non-native *A. thaliana* range. We also investigated population-specific groupings (bottom) such as if the population was collected before the year 2000, was collected non-native range, or is considered a relict (ice age refugia survivors).

The experimental gardens and populations included in this meta-analysis differ in many details of design and sampling, and this heterogeneity both bolsters the robustness of our species-wide mean adaptation lag estimate and provides an opportunity to examine how garden design and population attributes influence lag. Using the large sample size of our dataset (~30,000 observations), we conducted a series of exploratory re-analyses in which we re-fit the model to subsets defined by garden characteristics and accession traits (**Fig. 2C, Table S7**. Gardens located outside the native range of *A. thaliana* exhibited positive adaptation lags on average), consistent with the idea that non-native or invasive populations can confer novel environmental advantages that decouple populations from their historical climate optima (Callaway & Ridenour, 2004; Corbin & D’Antonio, 2010; Davidson et al., 2011). In contrast, several experimental design elements did not alter the sign of lag: whether plants were established as seeds versus seedlings or whether they were planted directly in the ground versus in pots, all subsets showed negative lag. However, pot-grown plants tended to show slightly less negative lags than ground-planted plants, suggesting that pots may buffer individuals from the full fitness costs of warming. Likewise, lag estimates from seeds versus seedling experiments were nearly indistinguishable, implying that temperature deviations in these studies did not differentially select on early life stage versus later establishment (Akiyama & Ågren, 2014; Schupp, 1995). Finally gardens that included water addition treatments produced lag estimates with notably broader confidence intervals than those without water addition, hinting at an interaction between temperature and moisture regimes that could modulate the expression of adaptation lag (Fréjaville et al., 2020; Morison & Morecroft, 2006). At the population level, most accession level groupings showed confidence intervals that crossed 0 and therefore were not contributing to significant estimates of lag. However, we observed suggestive trends; accessions collected outside the native range tended to show positive lags, relict populations from the southern glacial refugia had slightly less negative lags than non-relicts, and lag direction did not depend on the year of collection.

We then asked whether different populations had different levels of adaptation lag, how differences were distributed across space, and what are potential consequences of spatially structured adaptation lag. Our modeling approach and repeatability of populations being transplanted to multiple environments permitted us to generate genotype random effect estimates and therefore infer population-specific lag (*λ*_*i*_) (see **Supplemental Methods**). Ninety-three percent of populations exhibited lag less than 0°C, and we found some variation in each population’s lag, which ranged from ±3.15 °C around the mean (i.e., populations ranged from −5.06 °C to 1.24 °C) (**Fig. 2B**). The degree of lag was significantly correlated with latitude (*R*^*2*^ = 0.18, P = 3.54 × 10^−14^), where lower latitude populations had more negative lag values consistent with other measure of lag in plants across a latitudinal cline (Fréjaville et al., 2020).

Such extreme levels of adaptation lag in these high temperature regions could lead to a heightened fitness burden accumulating over years of unrealized fitness potential. Plant biomass is expected to suffer a loss that varies based on the future emission scenarios. In a conservative emission scenario (RCP4.5) biomass loss is predicted to range from 6.8-12%, while in a high emissions scenario (RCP 8.5) biomass loss ranges from 13.3-20.1% (Uribe et al., 2023). In *A. thaliana*, climate change has created relatively consistent warming across the range with 982 populations showing a mean annual-temperature deviation trend from TerraClim of +0.40 °C per decade 2000-2025 that is essentially identical across latitudinal bands (**Fig. 3**). Although such temperature deviations appear small, they translate into substantial fitness losses when mapped through the lag model. Rearranging the Gaussian fitness function, *decline(ΔT)=1-exp(−(ΔT*+*μ*_*λ*_ )^*2*^*/(2V*_*s*_ )), we estimate mean fitness declines of −24.6% [−48.0,-12.1], −25.2% [−46.5,-14.2], −27.2% [−49.0,-13.6], and −29.4% [−54.6,-12.9] under SSP1-2.6 (sustainability scenario), SSP2-4.5, SSP3-7.0, and SSP5-8.5 (fossil fuel centered scenario) respectively. These projected losses are in addition to the unmeasured fitness consequences of climate lag accumulated over the previous century. To better understand the potential impact of future emission scenarios on species’ adaptation and fitness, we extracted per-year climate projections (Thrasher et al., 2022) from global circulation models (GCMs) from present until 2050 that include four shared socioeconomic pathways, SSP1-2.6, SSP2-4.5, SSP3-7.0, and SSP5-8.5, spanning sustainable development, middle-of-the-road, regional rivalry, and fossil-fueled growth futures. NASA’s NEX-GDDP-CMIP6 ensemble mean deviations at 2050 vary from +2.59 °C under SSP1-2.6 to +3.21 °C under SSP5-8.5, with SSP2-4.5 (+2.85 °C) and SSP3-7.0 (+2.89 °C) in between. In addition, individual GCM × pixel year-to-year variability is ±0.5 °C, creating substantial deviations. Using the species-wide *λ*, mean per-year fitness at 2050 falls between −1.69 % (SSP1-2.6) and −2.23 % declines in fitness every year (SSP5-8.5), and a total reproductive deficit accumulated from 2000 to 2050 of −36.8% (SSP1-2.6) to −40.0% fitness (SSP5-8.5) (**Fig. 3B**).

**Figure 3.**
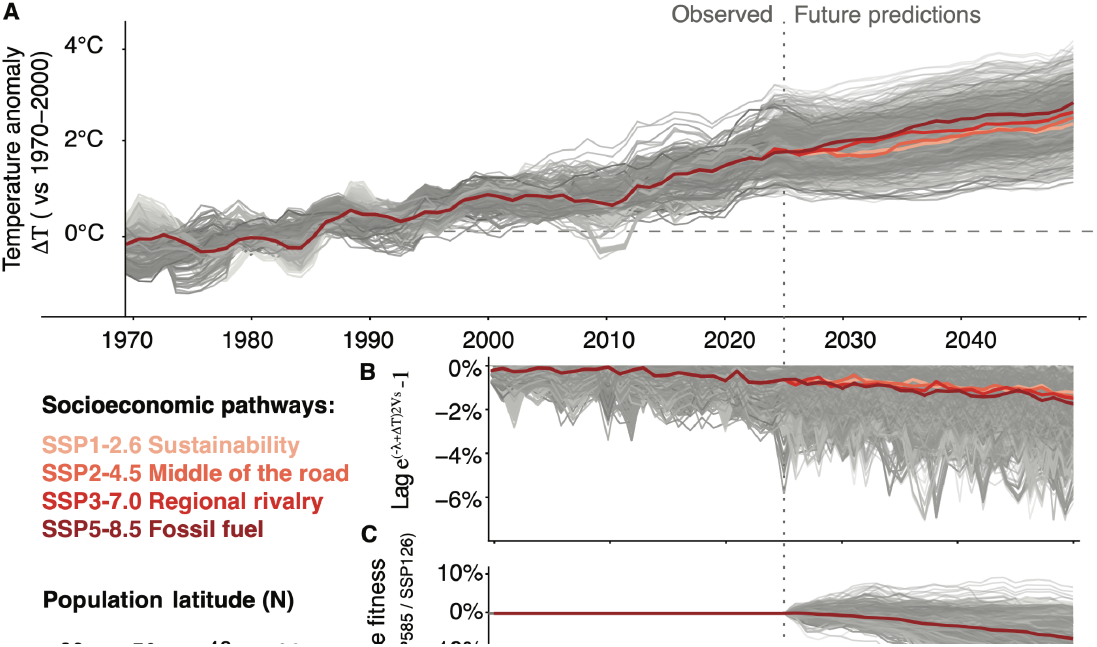
Projections of future adaptation lags under socio-economic global change scenarios. (A) Observed climate deviations 1970-2025 observed at each *Arabidopsis thaliana* population colored by latitude (source: TerraClim). Deviations are calculated for every year with respect to the historical 1970–2000 average (Worldclim). Future climate change predictions start in 2026 using 5 year moving averages. (B) Population fitness loss relative to the optimum estimated from transplants due to measured adaptation lag (λ = −1.91 °C) and ongoing temperature deviations (ΔT). In the future, multiple lines per location are plotted and thick lines represent the average across socioeconomic pathways scenarios. (C) Difference in cumulative fitness loss between a sustainable scenario (SSP1-2.6) and fossil fuel development (SSP5.8.5).

Finally, to directly attribute fitness deficit to future policy decisions, we compared the fitness in SSP1-2.6 (sustainability) and SSP5-8.5 (fossil fuel) scenarios. We found that failing to mitigate climate change based on Paris Agreement policies, erodes a median of −7.5% of fitness (10-90 percentile: −12.4 %, +0.8 %) by 2050 in addition to the already-large scenario of low emission/sustainability. While not the most ambitious sustainable scenario, SSP1-2.6 aims to limit global warming below 2 °C by 2100 by cutting emissions near zero by 2075 (SSP1-1.9 not included); although this scenario has already been rendered unlikely given the failing of policy commitments based on United Nations Environment Programme (UNEP) assessments and climate policy tracking (United Nations Environment Programme, 2025) (**Fig. 3C**). We cannot permit reaching the worst-case scenario, and we must re-uphold policies to at least an intermediate SSP2-4.5 scenario, which would nevertheless mean accepting a loss of −25.2% in fitness (**Fig. 3C** SSP4.5).

Our observations are consistent with a growing body of literature suggesting that adaptation lag may be more prevalent across plant species than previously appreciated. For example, in the Rock Mountains, a closely related mustard, *Boechera stricta*, had populations that appear unable to track earlier snowmelt through upslope migration, leading to maladaptation and motivating efforts toward genetic rescue (Anderson et al., 2025). Similarly, *Quercus lobata* populations in California have been shown to fall outside their historical climate niche (Browne et al., 2019). Applying our adaptation lag model, we extend these findings by quantitatively estimating evolutionary mismatch across species using publicly available transplant experiments (**Table 1**). Across 17 vascular plant species with diverse experimental designs, we were able to detect significant adaptation lags in four species using different fitness metrics. The perennial switchgrass (*Panicum virgatum*) showed −6.1 °C [−5.3, −6.8], the maritime pine (*Pinus pinaster*) at −6.0 °C, and the aforementioned Valley Oak (*Quercus lobata*) showed a clear lag but optimum could not be inferred because all gardens sit entirely on one flank of the thermal optimum, indicating at least a ≥ −6–8 °C maladaptation lag. Only one species, Atlantic white cedar (*Chamaecyparis thyoides*) yielded a reversed lag of +1.3 °C [+2.5, +0.1] (i.e. populations performed better in warmer environments), probably because only three gardens were assessed and the southernmost garden permitted winter survival independent on population of origin. We believe most species do not yet have sufficient data to yield significant results. A joint model across all species excluding *A. thaliana* with normalized fitness and taxon and study as random effect shows an average adaptation lag, although non-significant (−0.73°C [CI95% −4.36, +1.49], 75% of posterior distribution indicates lags). Even with statistical uncertainty and species-to-species heterogeneity, having such a lag across species would mean a massive impact on fitness of plant species worldwide in the next decades. Even under middle-of-the-road scenario, by mid-century any given year’s deviations will surpass the uncertainty threshold across such non-model species (SSP2-4.6, 2025 *ΔT*_*land*_ = 1.3°C), which we may estimate back-of-the envelope fitness loss of 1.8% fitness (CI95% 0.01% - 23.4%) (**Table S5**). The fossil-fuel-favoring socioeconomic scenario (SSP5-8.5, 2025 *ΔT*_*land*_ = +2.1°C) would imply a 3.5% fitness loss (0.06%-29.4%) (**Table S5**). Some species with the largest lags such as perennial bunchgrass, Pines, or Valley Oaks, are expected to have a cumulative fitness loss of 10 %, 22%, and 60%, respectively, in the optimistic scenario (SSP2-4.6) (**Table S5**). Because these impacts propagate across populations and across species, the compound fitness loss of ecosystems would be insurmountable (**Table S6**).

**Table 1.** Plant common-garden panel to re-fitting the climate-lag model. Species sorted by design power (N gardens × N source populations). Source coverage = fraction of the species’ native mean-annual-temperature range (from up to 1,500 GBIF occurrences per species, sampled on WorldClim v2.1 bio1) spanned by the study’s source populations. *d* range is the experimental *d*_temp_ = garden − source climate difference span. Gaussian-lag parameters *V*_*s*_ (niche width,C °C^2^) and (climate lag, °C — positive = adapted to cooler than current home) are from independent per-species MCMCglmm fits with no prior, using year-specific site climate. Values in **bold** have 95% credible intervals that exclude zero.

| Species | N gardens | N sources | Cov. (%) | $d_{\text{range}}$ ($^{\circ}\text{C}$) | $V_s$ [CI95] ($^{\circ}\text{C}^2$) | $\mu_{\lambda}$ [CI95] ($^{\circ}\text{C}$) |
| --- | --- | --- | --- | --- | --- | --- |
| *Arabidopsis thaliana* | 45 | 1,564 | 207% | 43.1 | 814.7 [716.8, 968.9] | **-1.9 [-2.9,-1.1]** |
| *Quercus lobata* | 2 | 658 | — | 4.6 | n/a ($\beta_2 > 0$) | **$\geq -6$ to $-8$** |
| *Panicum virgatum* | 10 | 592 | 67% | — | 109.7 [103.0, 116.9] | **-6.1 [-6.8, -5.3]** |
| *Fagus sylvatica* | 29 | 202 | 71% | 12.7 | 1,057 [335, 11,944] | -2.6 [-61.8,+6.3] |
| *Picea glauca* | 16 | 242 | 51% | 19.5 | 69.8 [52.1, 105.3] | +0.8 [-0.4, +1.8] |
| *Plantago lanceolata* | 16 | 38 | 78% | 29.6 | 353 [139, 5,763] | +4.3 [-3.8, +51] |
| *Juglans nigra* | 7 | 66 | 66% | 12.6 | 44 [23, 188] | -0.5 [-4.1,+4.9] |
| *Silene flos-cuculi* | 18 | 22 | 42% | 13.6 | 69 [22, 310] | +6.5 [-3.3,+35] |
| *Chamaecrista fasciculata* | 16 | 19 | 48% | 23.4 | 127 [50, 1,321] | +3.2 [-1.6, +8.3] |
| *Boechera stricta* | 5 | 52 | 22% | 4.4 | 8.4 [4.2, 47.5] | +1.5 [-0.0,+3.2] |
| *Lotus corniculatus* | 7 | 32 | 21% | 5.0 | 8 [3, 75] | +1.4 [+1.2, -5.9] |
| *Populus tremuloides* | 5 | 43 | 23% | 7.4 | 27 [15, 94] | +0.5 [-1.3, +2.7] |
| *Pinus pinaster* | 5 | 34 | 56% | 10.9 | 243.7 [104.9, 2,299.2] | **-6.0 [-51.7,-0.9]** |
| *Chamaecyparis thyoides* | 3 | 34 | 96% | 18.8 | 56 [42, 86] | **+1.3 [+0.1, +2.5]** |
| *Arabidopsis lyrata* | 7 | 12 | 33% | 18.5 | 127.3 [43.1, 1,120.4] | +3.7 [-7.5, +36.2] |
| *Pinus canariensis* | 4 | 21 | 88% | 11.9 | 155 [44, 1,466] | -5.0 [-28, +2.2] |
| *Erythranthe guttata* | 4 | 14 | 46% | 10.9 | 14 [7, 92] | -2.2 [-8.2, +0.1] |

**Notes.** *Q. lobata* $\mu_{\lambda}$ is a directional lower bound (gardens don't bracket the Gaussian peak); design power counts the 2 common gardens × 5 blocks each. *A. thaliana*'s source coverage exceeds 100% because the pipeline uses per-year TerraClimate annual T at source locations (with cold-year deviations down to $-11.7^{\circ}\text{C}$ ) while GBIF coverage is computed on the 1970–2000 climatology. **Species-level climate source.** GBIF retrieved 2026-04-20 via `rgbif::occ_search` (hasCoordinate = TRUE, hasGeospatialIssue = FALSE, limit 1,500). Bio1 overlay: WorldClim v2.1 10-arcmin.

Much work and literature on the ability of plants to adapt to climate change focuses on if there is enough standing genetic variation to enable adaptation (Barrett & Schluter, 2008). This analysis suggests that at least for a model system of relatively low conservation concern, standing genetic variation is not enough to adapt to climate change. Measuring adaptation lag across species will be essential for communicating with policy makers and predicting which species or populations are most at risk. While it is imperative to understand the extent of adaptation lag across species on Earth, in order to approach biology from a place of curiosity and wonder rather than urgency, the only true imperative is to slow the rate of global climate change.

## Additional information

## Acknowledgements

We thank Ruth Epstein, Miles Roberts, Ben Jin, Eddy Mendoza-Galindo, Meixi Lin, and other members of the MOILAB for their feedback on the manuscript.

## Author contribution

M.E.-A. and L.L. conceptualized the project, conducted analyses, and wrote the manuscript.

## Data availability

All scripts are available at Zenodo: <DOI add>.

## Supplementary Materials

### Methods and Supplemental Tables

#### Data collection

We searched for outdoor common garden experiments (73 studies total), many of which we had already documented in (Leventhal, Ruffley, and Exposito-Alonso 2025). Studies were published between 1993 and 2021 and mostly occurred in the northern hemisphere. We selected studies that used natural accessions (no mutant plants, mutation accumulation lines, or recombinant inbred lines), and most accessions were from the 1001 Genomes and RegMap, while some were wild accessions not in either resource. If a study was from an outdoor common garden and (1) contained more than one natural accessions and (2) recorded a component of fitness (survival, silique number, fruit length, seed count, seed weight, rosette diameter, inflorescence height, number of branches), we included that study in our analysis. Because it is naturalized globally, we included both accessions and gardens outside of the native range, although we have a column noting this in our data set to use a co-variate (see below). In total our data came from 10 studies spanning 48 gardens (**Table S1**) with experiments conducted between the years 2002 to 2021.

**Table S1.**
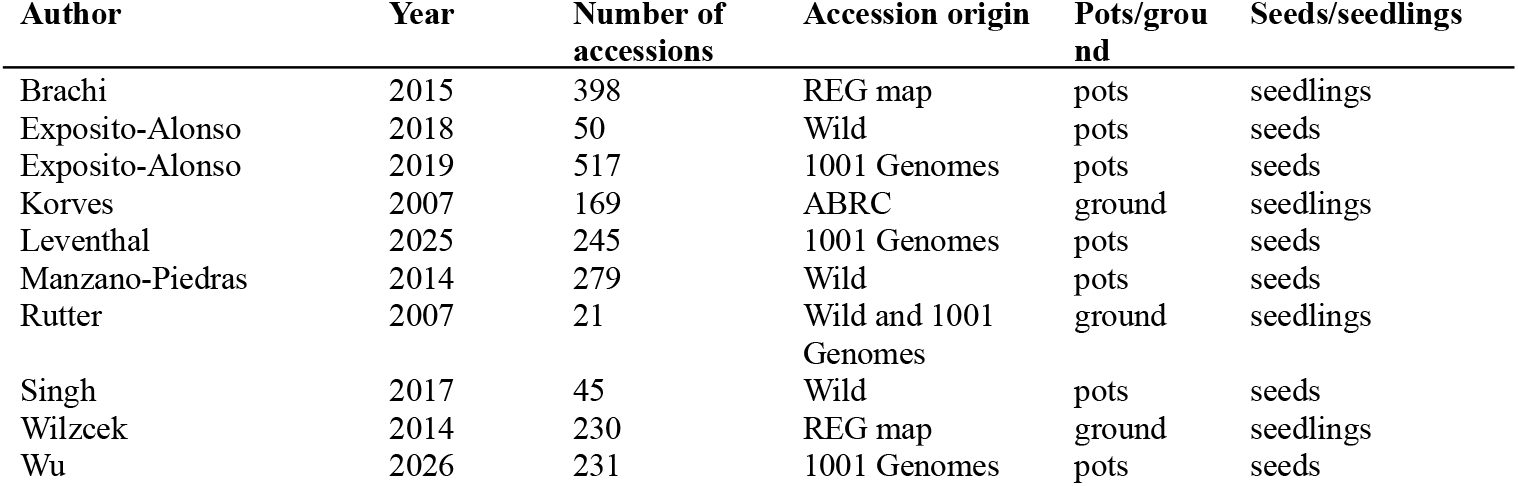
Summary of common garden experiments included in meta-analysis.

#### Fitness data

Across all fitness metrics, we had 78,856 observations across 1,655 distinct accessions. This includes a measurement for every accessions across the permutations of experiment (an individual publication), site (the experimental location), year (the year the garden was planted), season (the time of year the garden was planted, namely for Fall and Spring), and the treatment, which only applied to two of the studies. When we filtered the data to fecundity and lifetime fitness, there were 28,853 observations across 1614 distinct accessions. Some studies provided a measure of lifetime fitness (essentially survival × fecundity), but some studies would measure both survival and some reproductive output (e.g., fruit number or seed weight). To model lifetime fitness in the case of both survival and fecundity being provided, we modified these studies to have a lifetime fitness measurement by multiplying survival by fecundity. The fitness metrics that were used in the model for each study is included in **Table S4**.

The column is log-normalized within each garden 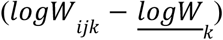. We apply a further within-garden min–max rescaling using the 1st–99th percentile as robust anchors, mapping every garden to the interval [0, 1]. This preserves rank order and makes *V*_*s*_ comparable across studies.

**Table S2.** Fitness metrics chosen from each common garden study.

| Study | Selected fitness metric | Rationale |
| --- | --- | --- |
| Brachi 2015 | Lifetime fitness | Pre-computed metric in original study (total length of mature fruits produced by each plant) |
| Exposito-Alonso 2018 | Survival × fruits | Mean number of rosettes (mean maximum number of rosettes per block) × Fruits (mean per reproductive plant) |
| Exposito-Alonso 2019 | Fitness | Pre-computed combined metric; each Density × Water combination is a distinct garden environment |
| Korves 2007 | Total seed mass × survival for winter survivors | Mean total seed mass per accession × Survival |
| Leventhal 2025 | Lifetime fitness | Survival (proportion, 0-1) × Fruits (mean per surviving plant); each precipitation treatment is a distinct garden |
| Manzano-Piedras 2014 | Rosettes × fruits | Pre-computed combined metric; single garden but genotypes overlap heavily with other studies |
| Rutter 2007 | Fruits | Pre-computed combined metric |
| Singh 2017 | Number of fruits | Pre-computed combined metric |
| Wilzcek 2014 | Seed number | Pre-computed fitness metric |
| Wu 2026 | Accession frequency | Integrates survival + fecundity + competition over 3 years × 31 gardens |

#### Climate data sources

We recorded the latitude and longitude of each experimental site from the published manuscripts, and if coordinates were not given, the closest possible location was selected. We collected the latitude and longitude for each accession from either the home database (1001 Genomes or RegBank) or in the case of wild accessions, from the respective manuscript. For the experimental locations climate data, we gather the year of the experiment and coordinates, and pulled the corresponding data from TerraClim. For the accessions’ climate data, we gathered the year the accession was collected from the 1001 Genomes or RegBank database or from the records of the authors in the case of wild accessions. If a range of years of collection was given for wild accessions, we selected the median year for those accessions. If the year of collection was not provided in the 1001 Genomes database or RegBank database, the median year of collection for all accessions was used. In the case that the collector of an accession was known but the year was not known, we used the median year for all accessions collected by that individual. Once every accession had a recorded year of collection, using TerraClim we pulled the climate for that year at the given accession’s coordinates. 4.7% (77/1617) of the accession were collected before 1958 (the earliest year of TerraClim data), then the year 1958 was used. From TerraClim, we selected monthly mean temperature (°C), monthly cumulative precipitation (mm), and the coefficient of variation in monthly temperature (higher values means more space in between rainfall events). We then calculated mean annual temperature (°C), annual precipitation accumulation (mm), and mean annual monthly coefficient of variation in precipitation.

##### Historical observations (1970–2025)

We extracted annual mean air temperature at every focal pixel from **TerraClimate** v1.5 (Abatzoglou et al. 2018), a 1/24° (≈ 4 km) global monthly reanalysis of minimum and maximum surface air temperature. For each pixel and calendar year we took the twelve monthly values of (*T*_*min*_ + *T*_*max*_)/2 and averaged them to a single annual mean *T*^−^_*y*_ covering 1970–2025. TerraClimate NetCDF files were downloaded in advance from the Climatology Lab and read locally with the **terra** R package.

##### Future projections (2026–2050)

Future air temperature came from the NASA NEX-GDDP-CMIP6 bias-corrected statistically downscaled ensemble (Thrasher et al. 2022), which provides daily mean near-surface air temperature (tas) on a 0.25° grid. We fetched all four standard Shared Socioeconomic Pathway scenarios spanning the emissions spectrum: SSP1-2.6 (sustainability, low emissions), SSP2-4.5 (middle of the road), SSP3-7.0 (regional rivalry, high emissions), and SSP5-8.5 (fossil-fuelled development). For every scenario we ran a five-GCM ensemble: ACCESS-CM2, EC-Earth3, MPI-ESM1-2-HR, MRI-ESM2-0, and UKESM1-0-LL, all at the r1i1p1f1 variant on each model’s native grid (except UKESM1-0-LL, which uses r1i1p1f2). NEX-GDDP daily files were retrieved one year at a time from the NASA Center for Climate Simulation THREDDS server (https://ds.nccs.nasa.gov/thredds/fileServer/AMES/NEX/GDDP-CMIP6/…), with incremental appends to a single cached CSV so the download is resumable. Daily values were averaged over each year to give an annual mean temperature per (pixel, scenario, GCM, year).

Cite NASA (10)

##### Per-GCM bias correction

TerraClimate (4 km, downscaled from CRU/WorldClim) and NEX-GDDP (25 km, bias-corrected against GMFD/ERA5) differ in absolute temperature scale by 0.5–2 °C in topographically complex regions. To make the two series directly stitchable we applied a per-GCM, per-pixel delta-method correction over the 2015–2025 overlap:

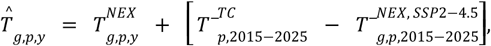

where *g* indexes GCM, *p* the 0.25° pixel, and *y* the year. The offset is computed from SSP2-4.5 only and reused across every scenario, because all four SSPs share almost identical forcings over 2015–2025 (we verified empirically that each GCM’s 2015–2025 mean varies by ≤ 0.3 °C across scenarios). Using a single-scenario overlap keeps every SSP on a common TerraClimate-anchored scale and preserves the relative differences between SSPs exactly.

##### Temperature deviations

Per-pixel deviations are expressed against a 1970–2000 TerraClimate baseline:

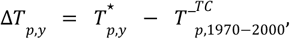

with 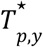 equal to TerraClimate’s annual mean for *y*≤2025 and to the bias-corrected NEX-GDDP value 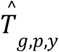 for *y*≥2026. TerraClimate observations 1970–2025 are identical across GCMs and scenarios and are replicated through every (scenario, GCM) channel so each projection trajectory is continuous from 1970 to 2050.

##### Visual smoothing

Annual values have ~0.4–0.5 °C year-to-year internal variability per GCM, which swamps the ~0.6 °C scenario divergence at 2050. For display we apply two sequential operations before plotting:

1. **GCM collapse**: for each (pixel, scenario, year) take the mean across the 5 GCMs. This removes model-internal variability from the display while preserving inter-scenario contrast.
2. **Five-year centred rolling mean** (`stats::filter` with `rep(1/5, 5)`, `sides = 2`) on each resulting per-(pixel × scenario) series. Smoothing is applied *after* the GCM collapse so it reduces noise without distorting ensemble means.

Per-scenario grand means across populations (the thick grey/black lines in Panel A) are computed from the GCM-collapsed series and smoothed with the same 5-year window.

#### Modeling adaptation lag

Here we define the notation used in the theoretical derivation of the lag model.

**Table S3.**
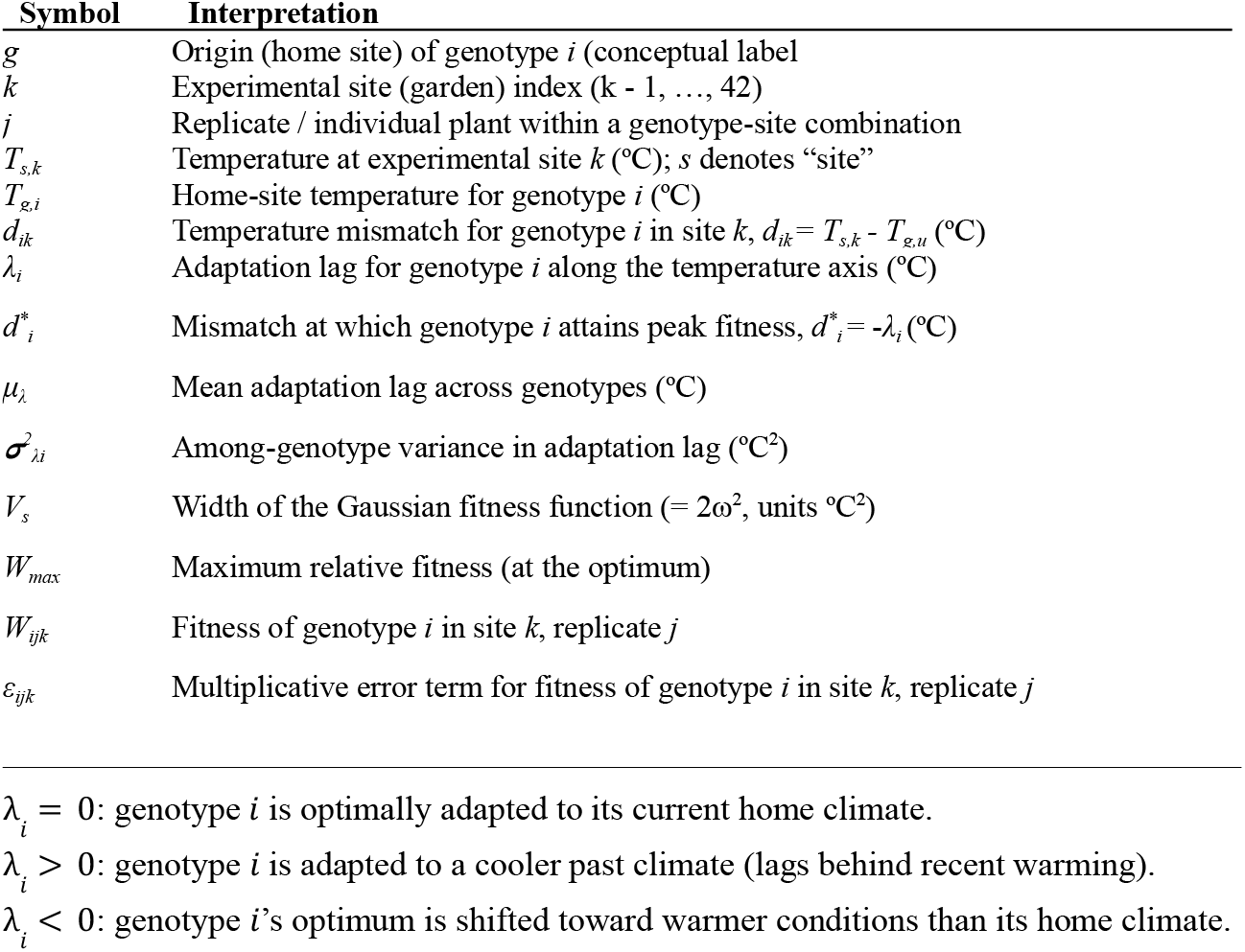
Mathematical symbols used in adaptation lag model.

| Symbol | Interpretation |
| --- | --- |
| $g$ | Origin (home site) of genotype $i$ (conceptual label) |
| $k$ | Experimental site (garden) index ($k = 1, \dots, 42$) |
| $j$ | Replicate / individual plant within a genotype-site combination |
| $T_{s,k}$ | Temperature at experimental site $k$ (°C); $s$ denotes “site” |
| $T_{g,i}$ | Home-site temperature for genotype $i$ (°C) |
| $d_{ik}$ | Temperature mismatch for genotype $i$ in site $k$, $d_{ik} = T_{s,k} - T_{g,i}$ (°C) |
| $\lambda_i$ | Adaptation lag for genotype $i$ along the temperature axis (°C) |
| $d_i^*$ | Mismatch at which genotype $i$ attains peak fitness, $d_i^* = -\lambda_i$ (°C) |
| $\mu_\lambda$ | Mean adaptation lag across genotypes (°C) |
| $\sigma_{\lambda i}^2$ | Among-genotype variance in adaptation lag (°C<sup>2</sup>) |
| $V_s$ | Width of the Gaussian fitness function ($= 2\omega^2$, units °C<sup>2</sup>) |
| $W_{max}$ | Maximum relative fitness (at the optimum) |
| $W_{ijk}$ | Fitness of genotype $i$ in site $k$, replicate $j$ |
| $\varepsilon_{ijk}$ | Multiplicative error term for fitness of genotype $i$ in site $k$, replicate $j$ |

$\lambda_i = 0$ : genotype $i$ is optimally adapted to its current home climate.
$\lambda_i > 0$ : genotype $i$ is adapted to a cooler past climate (lags behind recent warming).
$\lambda_i < 0$ : genotype $i$ 's optimum is shifted toward warmer conditions than its home climate.

A population is locally adapted when individuals perform best in a site whose climate matches their home climate. Under climate change, an **adaptation lag** arises when a genotype is still tuned to a *past* optimum: it reaches peak fitness not at sites matching its current home climate *T*_*g*_, but at sites whose climate resembles *T*_*g*_ − *λ*_*i*_ (a cooler past state) (visualized with raw data in **Fig. S2**). We model relative fitness as a Gaussian function of the difference between the site climate *T*_*s*_ and the genotype’s home climate *T*_*g*_ : *W*_ijk_ = *W*_max_ · exp[−(*T*_s,k_ − (*T*_g,*i*_ − λ_*i*_))^2^ / *V*_s_] · ε_ijk_.

To account for variations in genotype-level lag variation, we denote per genotype lag to *λ*_*i*_.

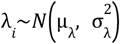

μ_*λ*_: mean lag across all genotypes. σ_*λ*_: standing genetic variation in thermal optimum.

In this Gaussian fitness function, *W*_*max*_ controls the height of the curve (maximum relative fitness), *V*_*s*_ controls its width (how quickly fitness declines as mismatch increases), and the term in the numerator of the exponent, (*T*_*s,k*_ – (*T*_*g,i*_ − *λ*_*i*_)) ^2^, determines where the peak falls along the temperature axis. The lag parameter *λ*_*i*_ shifts the optimum along this axis, so genotypes with different *λ*_*i*_ values have different peak climates.

For algebraic convenience, we re-express the model in terms of temperature mismatch. If we define the climate distance as *d*_*ik*_ = *T*_*s,k*_ − *T*_*g,i*_, the model becomes:

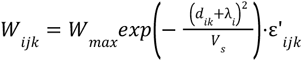

When we transform this model to a log scale our full Gaussian model becomes:

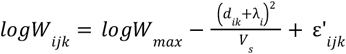

where ε’_*ijk*_ is the residual error on the log scale. For notational simplicity, we write this term as ε in the derivations below.

The fitness peak in *d*-space is at *d*^*^ =− *λ*_*i*_: a lagged genotype (*λ*_*i*_ > 0) reaches peak fitness at sites *cooler* than its home (*d* < 0). Because the log-Gaussian is a downward-opening parabola in *d*_*ik*_, we can expand (*d*_*ik*_ + *λ*_*i*_)^2^ and rewrite the model as a quadratic function of mismatch:

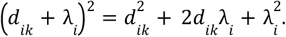

Distributing − 1/*V* gives

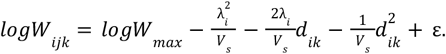

We can recognize this as

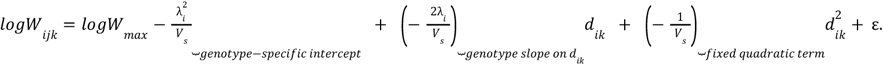

This suggests a random-slope mixed model with a fixed quadratic term:

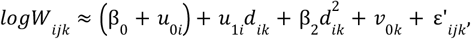

where *u*_0*i*_ and *u*_1*i*_ are genotype-specific random intercepts and slopes, β_2_ is a fixed quadratic effect shared across genotypes, and *v*_0*k*_ is a garden random intercept absorbing unmeasured garden-level variation.

**Table S4.**
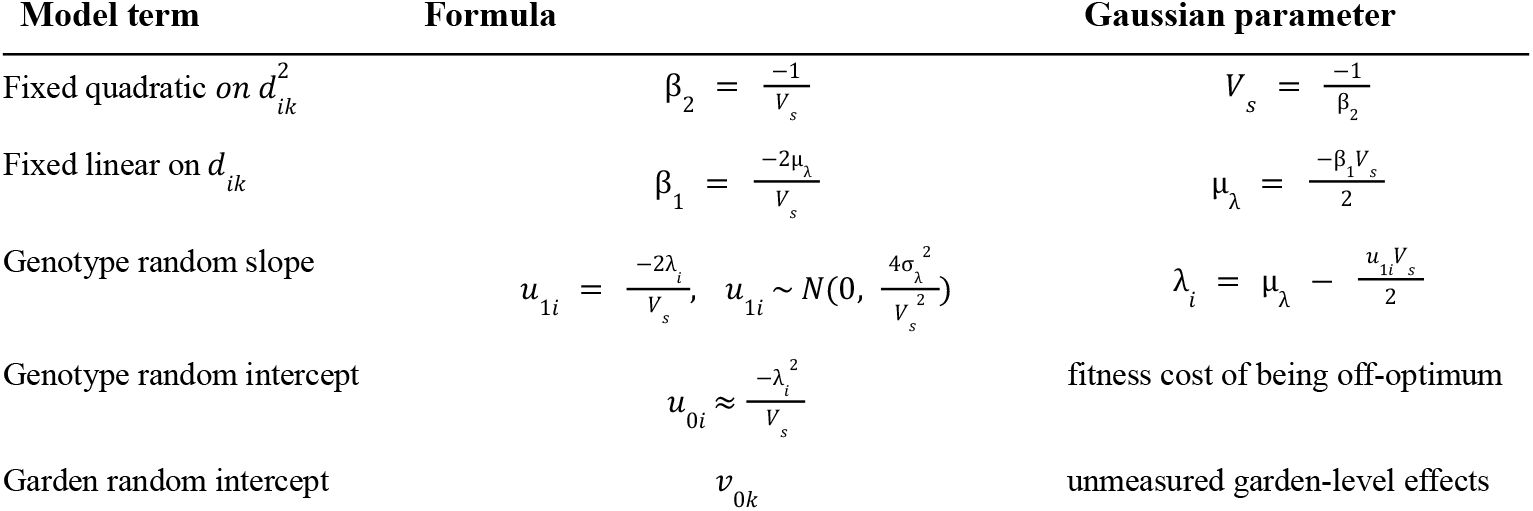
Mapping between mixed-model parameters and Gaussian lag parameters.

Under this reparameterization, the data identify the lag distribution through the fixed and random effects in the quadratic model. Genotype random slopes are informed by the pattern of fitness across the range of *d*_*ik*_ values observed for each genotype, and study-level intercepts absorb differences in absolute fitness among gardens.

Specifically, the model allows us to estimate (1) The **mean lag** μ_*λ*_, which is identified by the fixed linear slope β_1_ relating log fitness to temperature mismatch *d*_*ik*_. This parameter describes the average displacement of the fitness optimum along the temperature axis across all genotypes, (2) the **standard deviation of lag** σ_*λ*_, which is identified from the variance of the genotype-specific random slopes *u*_1*i*_ (this variance quantifies how much genotypes differ from one another in their lag, i.e. the spread of thermal optima around the mean), and (3) the **individual deviations** *λ*_*i*_ − μ_*λ*_ for each genotype, which are obtained as the best linear unbiased predictors (BLUPs) of the random slopes and are centered at zero by definition of the random-effects structure. These deviations show which genotypes lag more or less than the average and by how much. We do not attempt to estimate a separate *W*_*max*_ for each genotype, because fitness is rescaled within gardens and garden-level intercepts *v*_0*k*_ absorb differences in absolute fitness among experiments.

#### Fitness projections into future climates

##### Gaussian lag model

Per-year relative fitness is drawn from the Gaussian-selection reparameterisation of the Leventhal & Expósito-Alonso (2026, in prep.) MCMCglmm fit (`fit_main_10_to_10_year_fixed.rds`):

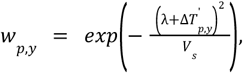

where *V*_*s*_ is the species-level selection width (posterior median 824.2) and *λ* is the per-population lag. We re-anchored each trajectory at year 2000 so 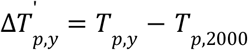. Year 2000 is the end of the 1970–2000 climatology baseline, so anchoring there keeps the fitness reference (0 %) conceptually consistent with the temperature-deviation baseline while still letting the trajectory accumulate the historical warming observed over 2000–2025.

Relative fitness at warming Δ*T* above today:

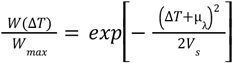

Fractional decline already present at current climate (“baseline” lag cost):

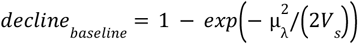

Fractional decline at future warming Δ*T*:

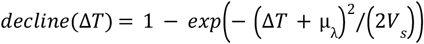

##### Two lag variants

We ran the same pipeline with two choices of *λ*:

1. **Population-specific lag (***λ*_*i*_). Each population uses its own posterior-median random-slope BLUP *λ*_*i*_. The across-population spread in *λ*_*i*_ captures biological variation in adaptation lag.
2. **Species-average lag (**μ_*λ*_). All populations share the species mean μ_*λ*_ (posterior median +1.96 °C), computed as μ_*λ*_ =− β_1_ *V*_*s*_ /2 from the fit’s fixed effects. Using μ_*λ*_ removes biological variation from the display so any remaining across-population spread reflects only differences in home climate (through *T*_*p*,2000_) and scenario/GCM uncertainty. Figure 3 uses the species-average formulation.

##### Per-year relative fitness

For every (population, scenario, year) we computed *w*_*p,y*_ using the species-wide μ_*λ*_ draw attached to that scenario’s GCM (see *Posterior sampling*). We then averaged across the 5 GCMs per (population, scenario, year) and smoothed the resulting series with a 5-year centred rolling mean, identical to the climate-data treatment. The plotted y-value is *w*_*p,y*_ − 1 expressed as a percentage, so 0 % corresponds to the undisturbed Gaussian optimum (*λ* = 0, Δ*T*^’^ = 0) and negative values indicate a yearly fitness deficit relative to that optimum.

##### Cumulative fitness shortfall under the worst-case scenario

Cumulative fitness from the 2000 anchor compounds the per-year values multiplicatively:

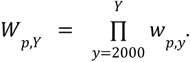

To compare scenarios we paired each population × GCM combination and took the ratio of cumulative fitness under SSP5-8.5 (fossil-fuelled, worst-case) relative to SSP1-2.6 (sustainability, best-case):

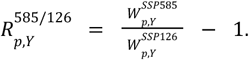

*R* < 0 indicates that SSP5-8.5 imposes a greater cumulative fitness cost than SSP1-2.6; *R* > 0 indicates the rare case where a population’s optimum sits above its current climate and would benefit from faster warming (*λ*_*i*_ < 0). *λ* is *shared* between the two scenario sides within each (population × GCM) pair, so the ratio isolates scenario effects rather than *λ*-posterior noise. The ratio series is smoothed with the same 5-year rolling mean.

##### Posterior sampling of the lag

We propagate posterior uncertainty on the lag by drawing from a Normal approximation to its 95 % credible interval:

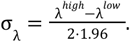

For the species-average variant used in Figure 3, μ_*λ*_ is a single scalar; we draw one posterior sample per GCM to simulate the dispersion expected due to uncertainty. Because we have many populations with variable climate trajectories, and we have multiple GCM scenarios, this gives independent samples of μ_*λ*_ capturing a natural posterior fan that is more computationally efficient without the need for time-consuming replicate loops. All populations under a given GCM share the same μ_*λ*_ draw, so the SSP5-8.5-vs-SSP1-2.6 ratio stays well-behaved under tail samples (if the two scenario sides used independent draws, extreme lag values in one direction would send the ratio to implausible values). *V*_*s*_ is not resampled and is fixed at its posterior median.

##### Average species loss

Taking *A. thaliana* aside, using a model of non-Arabidopsis species (also excluding *Quercus lobata*) we can make predictions of fitness loss in 2050.

**Table S4.**
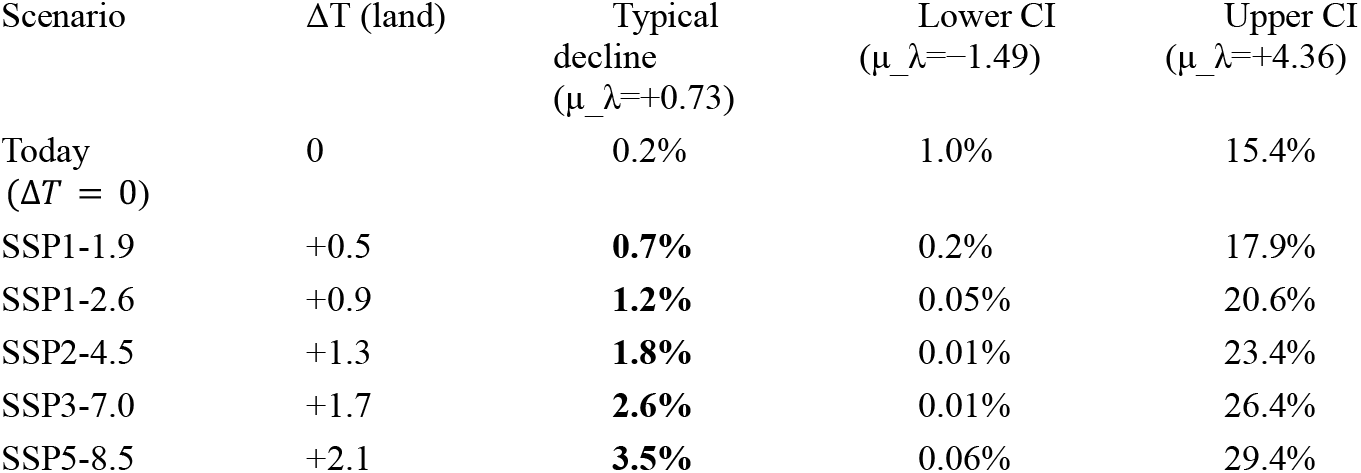
Potential fitness loss in different climate change socioeconomic scenarios. **Parameters:** μ_λ_ =+ 0. 73 [−1.49, +4.36] °C; *V*_*s*_ = 112 [60, 324] °C^2^ (σ_s ≈ 10.6 °C).

| Scenario | $\Delta T$ (land) | Typical decline<br>($\mu_{\lambda}=+0.73$) | Lower CI<br>($\mu_{\lambda}=-1.49$) | Upper CI<br>($\mu_{\lambda}=+4.36$) |
| --- | --- | --- | --- | --- |
| Today<br>($\Delta T = 0$) | 0 | 0.2% | 1.0% | 15.4% |
| SSP1-1.9 | +0.5 | **0.7%** | 0.2% | 17.9% |
| SSP1-2.6 | +0.9 | **1.2%** | 0.05% | 20.6% |
| SSP2-4.5 | +1.3 | **1.8%** | 0.01% | 23.4% |
| SSP3-7.0 | +1.7 | **2.6%** | 0.01% | 26.4% |
| SSP5-8.5 | +2.1 | **3.5%** | 0.06% | 29.4% |

At the Model G’ typical-plant level, 2050 land-warming causes a monotone fitness decline of **~0.7–3.5%** under the median posterior, with the upper CI reaching **~30%** if the true lag is near +4 °C. Per-decade rate under SSP2-4.5 ≈ **0.45 percentage points per decade** at the aggregate typical-plant level.

**Table S5.** Potential fitness losses of each species. Assuming correct inference of lag of course, and providing

| Species | $\mu_{\lambda}$<br>(°C) | $V_s$<br>(°C<sup>2</sup>) | Baseline today | SSP1-1.9 | SSP1-2.6 | SSP2-4.5 | SSP3-7.0 | SSP5-8.5 |
| --- | --- | --- | --- | --- | --- | --- | --- | --- |
| *Arabidoss thaliana* | +1.91 | 815 | 0.2% | 0.4% | 0.5% | 0.6% | 0.8% | 1.0% |
| *Quercus lobata* * | +6.00 | ~30 | **45%** | **51%** | **55%** | **59%** | **63%** | **67%** |
| *Panicum virgatum* | +6.10 | 110 | **16%** | **18%** | **20%** | **22%** | **24%** | **26%** |
| *Fagus sylvatica* | +2.60 | 1,057 | 0.3% | 0.5% | 0.6% | 0.7% | 0.9% | 1.0% |
| *Picea glauca* | −0.80 | 70 | 0.5% | 0.1% | 0.0% | 0.2% | 0.6% | 1.2% |
| *Plantago lanceolata* | −4.30 | 353 | 2.6% | 2.0% | 1.6% | 1.3% | 1.0% | 0.7% |
| *Juglans nigra* | +0.50 | 44 | 0.3% | 1.1% | 2.2% | 3.6% | 5.4% | 7.4% |
| *Silene flos cuculi* | −6.50 | 69 | **26%** | **23%** | **20%** | **18%** | **15%** | **13%** |
| *Chamaeristasta fasciculata* | −3.20 | 127 | 4.0% | 2.8% | 2.1% | 1.4% | 0.9% | 0.5% |
| *Boechera stricta* | −1.50 | 8 | **13%** | 6.1% | 2.2% | 0.2% | 0.2% | 2.2% |
| *Lotus corniculatus* | −1.40 | 8 | **12%** | 4.9% | 1.6% | 0.1% | 0.6% | 3.0% |
| *Populus tremuloides* | −0.50 | 27 | 0.5% | 0.0% | 0.3% | 1.2% | 2.6% | 4.6% |
| *Pinus pinaster* | +6.00 | 244 | 7.1% | 8.3% | 9.3% | **10.3%** | **11.4%** | **12.6%** |
| *Chamaecyparis thyoides* | −1.30 | 56 | 1.5% | 0.6% | 0.1% | 0.0% | 0.1% | 0.6% |
| *A. lyrata* ssp. *petraea* | −3.70 | 127 | 5.2% | 4.0% | 3.0% | 2.2% | 1.6% | 1.0% |
| *Pinus canariensis* | +5.00 | 155 | 7.7% | 9.3% | **10.6%** | **12.0%** | **13.5%** | **15.0%** |
| *Erythranthe guttata* | +2.20 | 14 | **16%** | **23%** | **29%** | **35%** | **42%** | **48%** |

\* *Quercus lobata* $V_s$ is a placeholder (our fit gave $\beta_2 > 0$ , not identifiable); value assumes Browne's among-source summer-Tmax SD $\approx 2.5$ – $3.5$ °C and $\mu_{\lambda} \approx +6$ °C lower bound.

**Table S6.** Cumulative aggregate of fitness decline (2050 land warming)

Summing across the 17-species panel:
| Scenario | Cumulative decline ($\Sigma$ species, %) | Mean per-species decline (%) |
| --- | --- | --- |
| SSP1-1.9 | **154** | 9.1 |
| SSP1-2.6 | **158** | 9.3 |
| SSP2-4.5 | **168** | 9.9 |
| SSP3-7.0 | **184** | 10.8 |
| SSP5-8.5 | **205** | **12.1** |

**Table S7.** Curiosity testing lag measurements based on accession or garden characteristics.

| Scenario-type | Scenario | mean $\lambda$ [95% CI] | Observation size |
| --- | --- | --- | --- |
| Experimental | Garden located in non-native range | 1.8[0.5:2.9] | 7295 |
| Experimental | Garden located in native range | -1.8[-2.7:-0.9] | 20372 |
| Experimental | Experiment planted in ground | -6.1[-29.6:-1.3] | 1199 |
| Experimental | Experiment planted in pots | -1.8[-2.9:-0.9] | 26468 |
| Experimental | Plants started outside as seedlings | -1.6[-4.2:0] | 2460 |
| Experimental | Plants started outside as seeds | -2[-3.2:-0.8] | 25207 |
| Experimental | Water was added to experimental design | -38.9[-159.5:82.2] | 6879 |
| Experimental | No water was added to experimental design | -1.9[-3:-0.8] | 23548 |
| Population | Mediterranean population | -11.1[-25.2:-1.5] | 13042 |
| Population | Non-mediterranean population | 82.5[-242.1:299.3] | 14625 |
| Population | Population collected before year 2000 | -0.8[-2.7:0.5] | 9887 |
| Population | Populations collected after year 2000 | -2.7[-4.4:-1.3] | 17780 |
| Population | Population present in 2 or more gardens | -2.5[-3.7:-1.5] | 26192 |
| Population | Population present in only 1 garden | 0.6[-6.8:5.9] | 1475 |
| Population | Population from non-native range | 2.8[-6.1:21.1] | 607 |
| Population | Population from native range | -1[-2:-0.2] | 27060 |
| Population | Populations are southern relicts | -1.9[-5:0.4] | 1134 |
| Population | Populations are not southern relicts | -2.4[-3.8:-1.4] | 26533 |

**Table S8.**
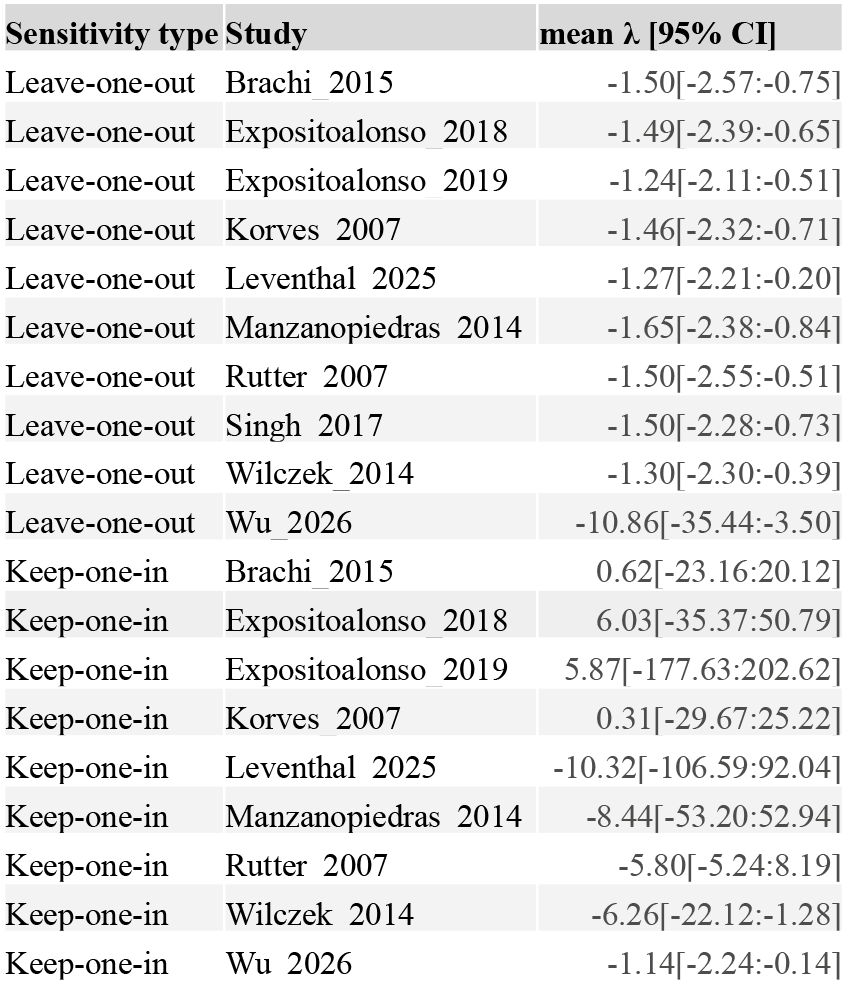
Sensitivity testing, leave one out and keep one in results.

| Sensitivity type | Study | mean $\lambda$ [95% CI] |
| --- | --- | --- |
| Leave-one-out | Brachi_2015 | -1.50[-2.57:-0.75] |
| Leave-one-out | Expositoalonso_2018 | -1.49[-2.39:-0.65] |
| Leave-one-out | Expositoalonso_2019 | -1.24[-2.11:-0.51] |
| Leave-one-out | Korves_2007 | -1.46[-2.32:-0.71] |
| Leave-one-out | Leventhal_2025 | -1.27[-2.21:-0.20] |
| Leave-one-out | Manzanopiedras_2014 | -1.65[-2.38:-0.84] |
| Leave-one-out | Rutter_2007 | -1.50[-2.55:-0.51] |
| Leave-one-out | Singh_2017 | -1.50[-2.28:-0.73] |
| Leave-one-out | Wilczek_2014 | -1.30[-2.30:-0.39] |
| Leave-one-out | Wu_2026 | -10.86[-35.44:-3.50] |
| Keep-one-in | Brachi_2015 | 0.62[-23.16:20.12] |
| Keep-one-in | Expositoalonso_2018 | 6.03[-35.37:50.79] |
| Keep-one-in | Expositoalonso_2019 | 5.87[-177.63:202.62] |
| Keep-one-in | Korves_2007 | 0.31[-29.67:25.22] |
| Keep-one-in | Leventhal_2025 | -10.32[-106.59:92.04] |
| Keep-one-in | Manzanopiedras_2014 | -8.44[-53.20:52.94] |
| Keep-one-in | Rutter_2007 | -5.80[-5.24:8.19] |
| Keep-one-in | Wilczek_2014 | -6.26[-22.12:-1.28] |
| Keep-one-in | Wu_2026 | -1.14[-2.24:-0.14] |

### Supplemental Figures

**Figure S1.**
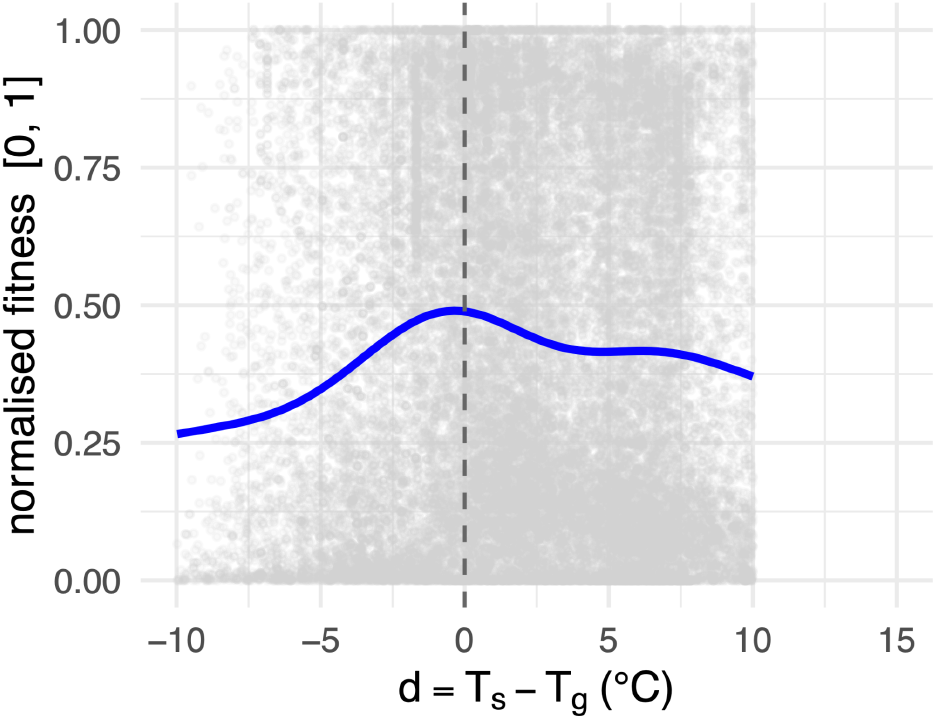
Relationship between fitness and annual temperature mismatch between experimental and genotype home site. Each point shows the normalised fitness of a population in a common garden as a function of d-space. The blue curve shows a smoothed fit (GAM) with 95% point-wise uncertainty omitted for clearing viewing of the raw fitness points. The vertical dashed line means no mismatch in temperature (d = 0),

**Figure S2.**
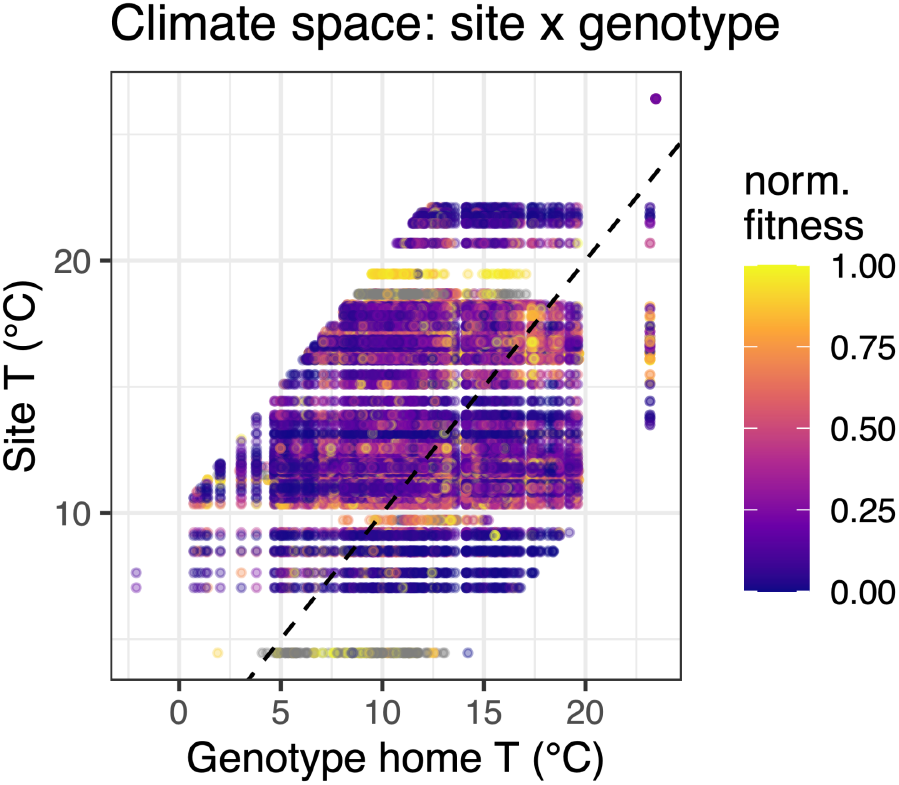
Climate space of common garden sites and genotypes. Each point represents a genotype-by-site combination, plotted by the long-term mean temperature at the genotype’s home site (x-axis) and the common garden site (y-axis), with point color indicating normalized fitness. The dashed 1-1 line denotes perfect temperature matching between home and garden climates, illustrating that some of the genotypes we used are well-matched while others are not well-matched to their experimental site.

**Figure S3.**
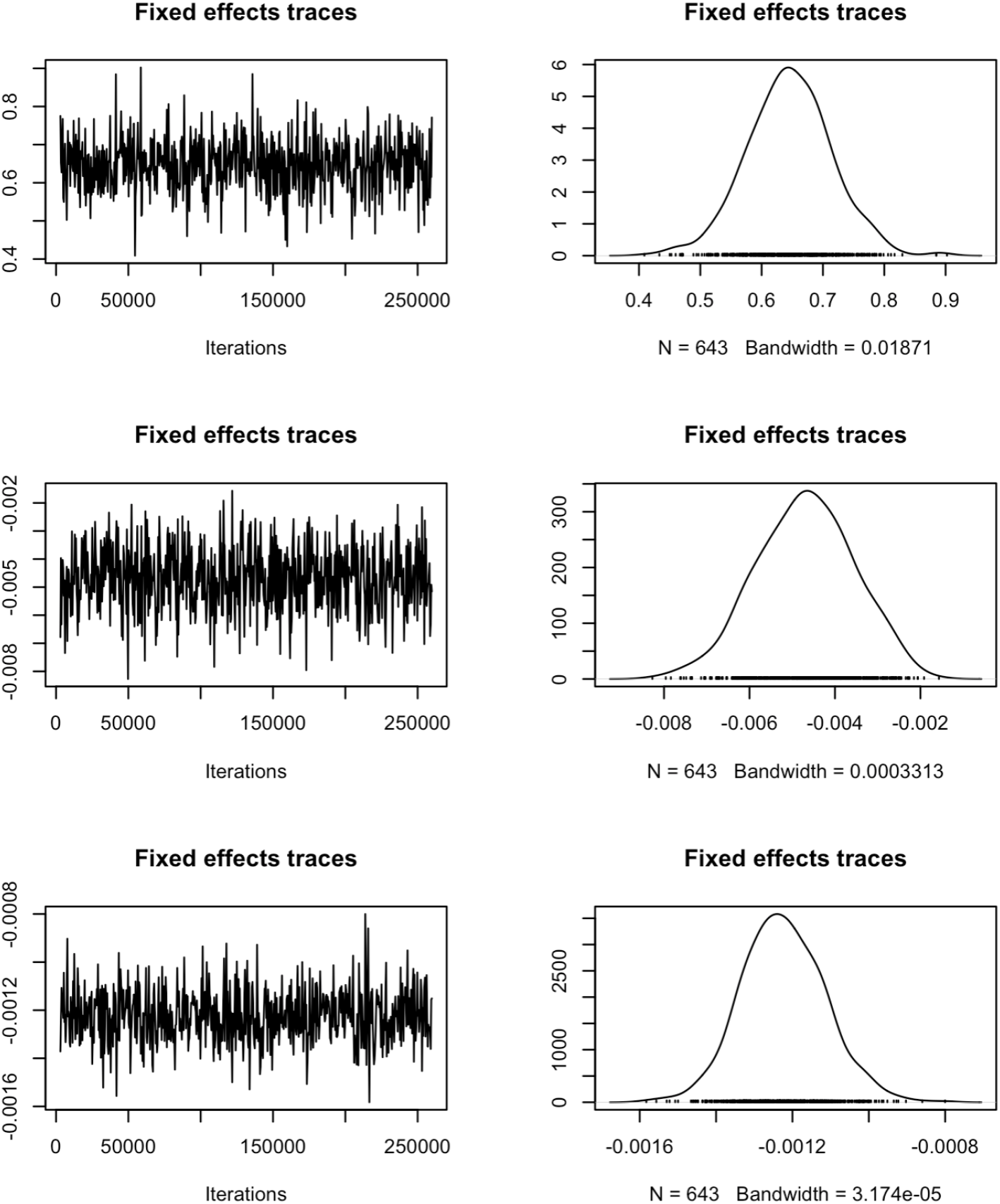
Model diagnostics for fixed effects in the adaptation-lag model. Trace plots and marginal posterior densities for fixed effects indicate good mixing and approximate convergence of the MCMC chains.

**Figure S4.**
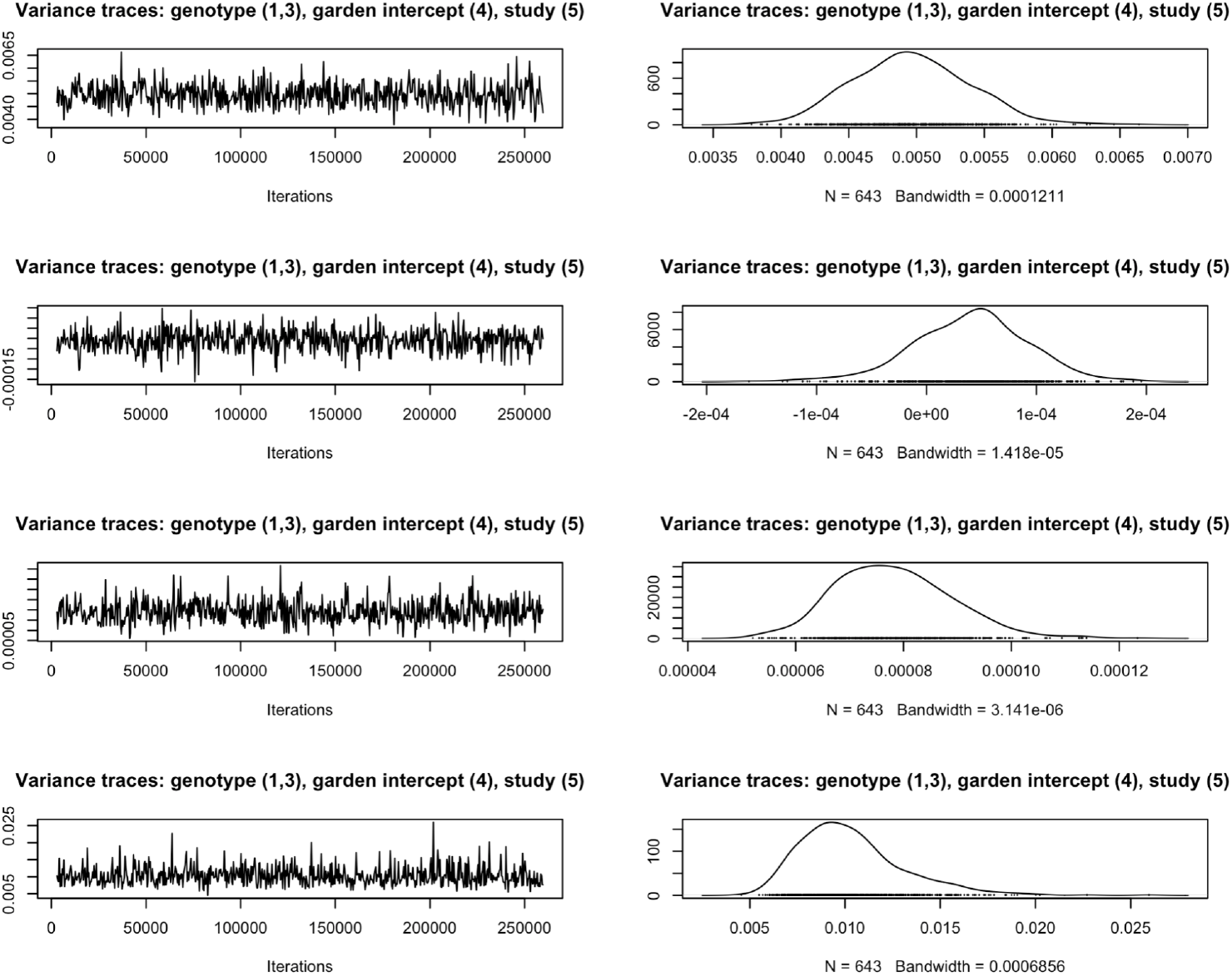
MCMC diagnostics for variance components in the adaptation-lag model. Trace plots and marginal posterior densities for the genotype, garden (site) intercept, and study-level random effect variances indicate good mixing and approximate convergence of the MCMC chains.

**Figure S5.**
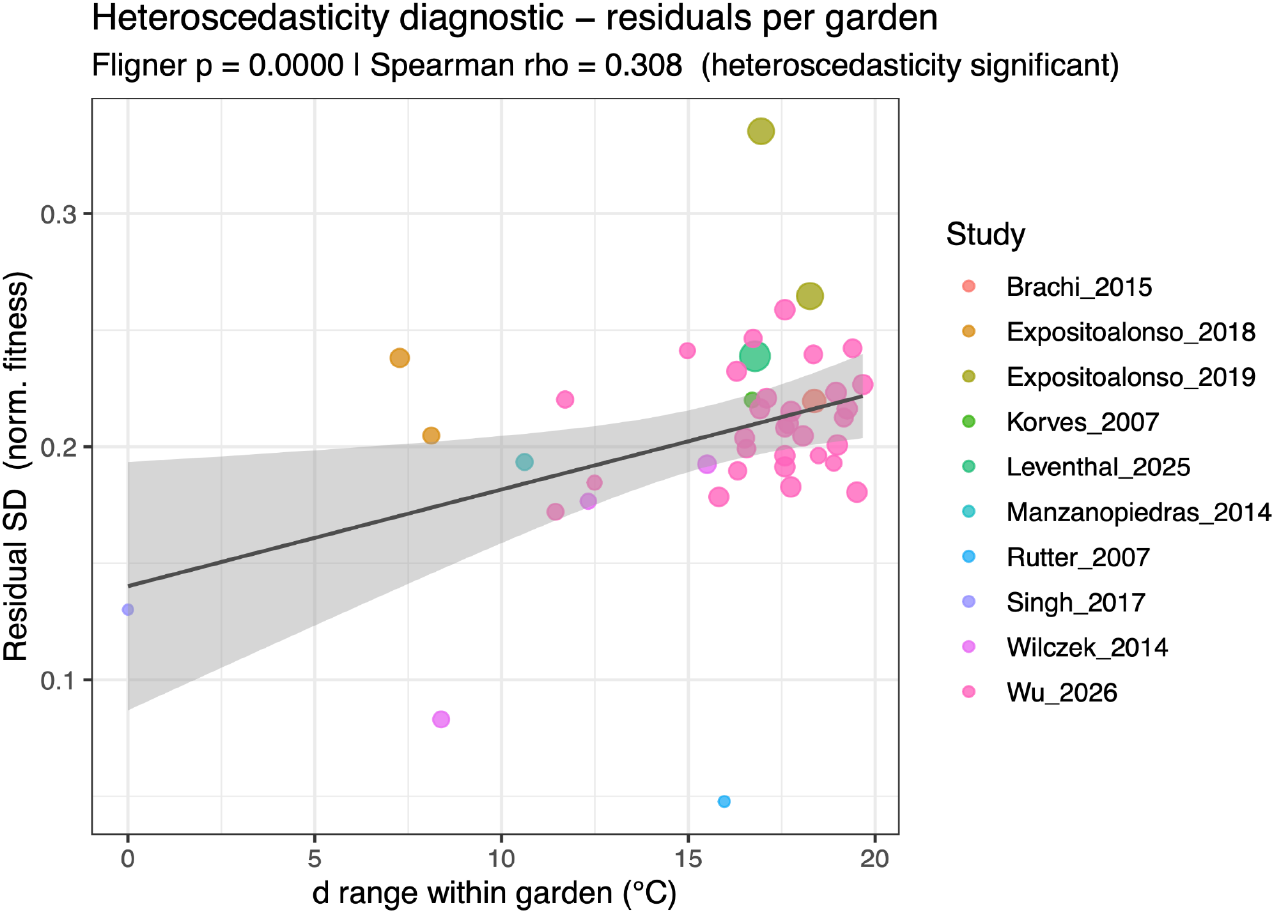
Heteroscedasticity diagnostics for the adaptation-lag model. Each point shows the standard deviation of model residuals (normalized fitness) with a garden plotted against the range of temperature mismatch *d* experienced in that garden, with colors indicating study identity and a fitted linear trend line illustrating increasing residual variance with increasing *d* range.

**Figure S6.**
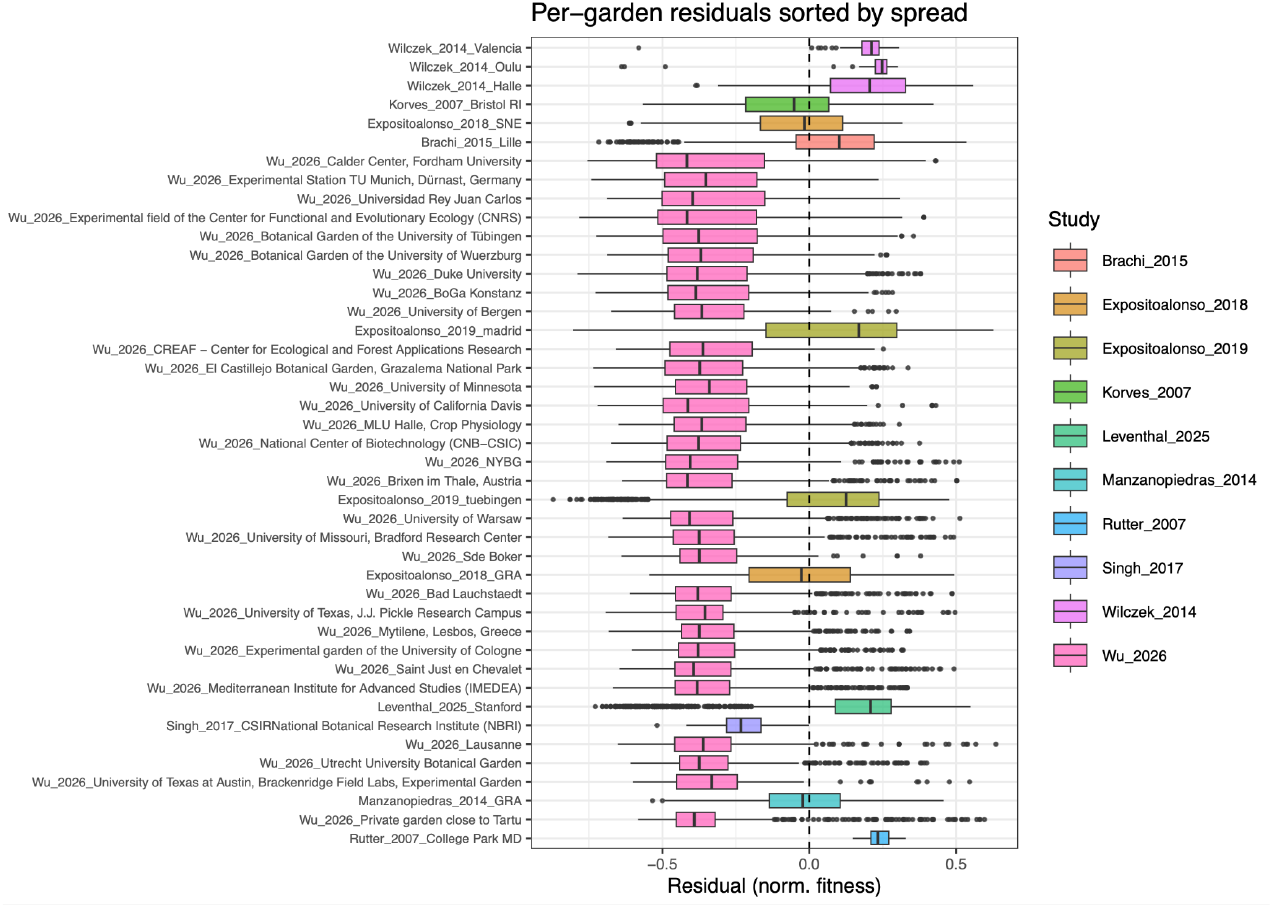
Heteroscedasticity diagnostics showing per-garden distribution of residuals from the adaptation-lag model. Boxplots show the distribution of normalized fitness residuals for each common garden, ordered by residual spread, with colors indicating study identity and the dashed vertical line marking zero residual (perfect model fit).

**Figure S7.**
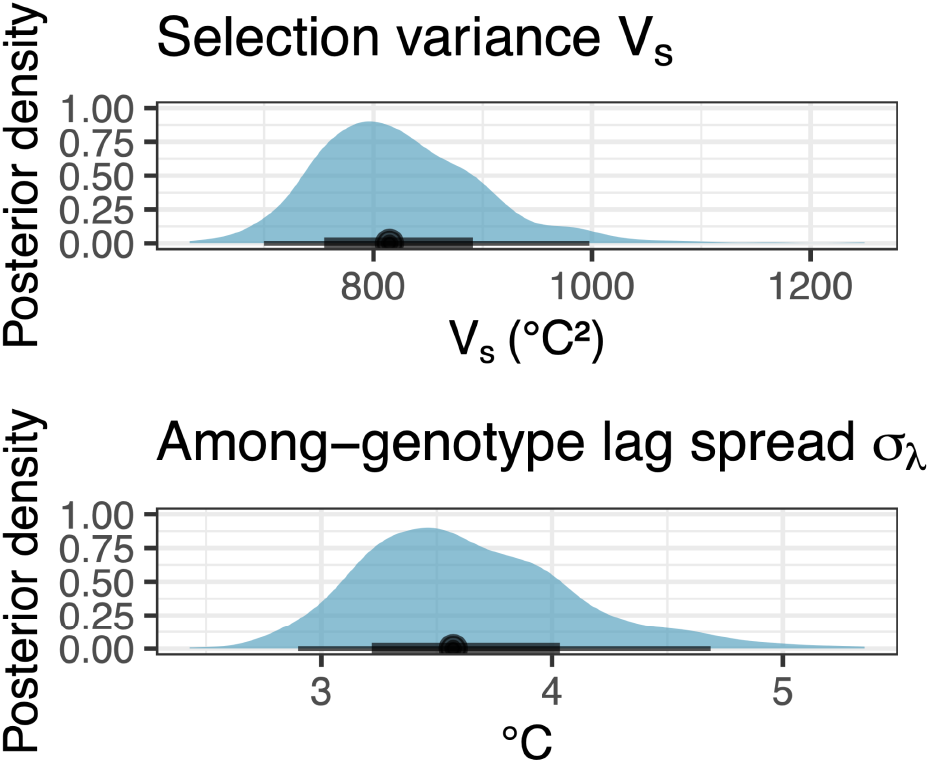
Posterior distributions for key adaptation-lag parameters. Shaded densities show the posterior for the selection and variance *V*_*s*_ (top; units °C^2^) and the among genotype lag spread *σ*_*λ*_ (bottom; units °C^2^), with horizontal bars indicating central credible intervals and points marking posterior medians.

